# Fast volumetric imaging with line-scan confocal microscopy by an electro-tunable lens

**DOI:** 10.1101/2021.12.01.470673

**Authors:** Khuong Duy Mac, Muhammad Mohsin Qureshi, Myeongsu Na, Sunghoe Chang, Hyuk-Sang Kwon, Tae Joong Eom, Hyunsoo Shawn Je, Young Ro Kim, Euiheon Chung

**Affiliations:** Department of Biomedical Science and Engineering, Gwangju Institute of Science and Technology, Gwangju 61005, Korea; Division of Biophysics and Bioimaging, Princess Margaret Cancer Centre, Toronto, ON, Canada; Department of Physiology and Biomedical Sciences, Seoul National University College of Medicine, 103 Daehak-ro, Jongno-gu, 03080 Seoul, South Korea; Neuroscience Research Institute, Seoul National University College of Medicine, 103 Daehak-ro, Jongno-gu, 03080 Seoul, South Korea; Advanced Photonics Research Institute, Gwangju Institute of Science and Technology, Gwangju 61005, South Korea; Signature Program in Neuroscience and Behavioural Disorders, Duke-National University of Singapore (NUS) Medical School, 8 College Road, Singapore, 169857, Singapore; Advanced Bioimaging Center, Academia, Ngee Ann Kongsi Discovery Tower Level 10, 20 College Road, Singapore 169855; Athinoula A. Martinos Center for Biomedical Imaging, Massachusetts General Hospital, Charlestown, Massachusetts 02129, USA; Department of Radiology, Harvard Medical School, Boston, Massachusetts 02115, USA; AI Graduate School, Gwangju Institute of Science and Technology, Gwangju 61005, Korea

## Abstract

In microscopic imaging of biological tissues, particularly real-time visualization of neuronal activities, rapid acquisition of volumetric images poses a prominent challenge. Typically, two-dimensional (2D) microscopy can be devised into an imaging system with 3D capability using any varifocal lens. Despite the conceptual simplicity, such an upgrade yet requires additional, complicated device components and suffers a reduced acquisition rate, which is critical to document neuronal dynamics properly. In this study, we implemented an electro-tunable lens (ETL) in the line-scan confocal microscopy, enabling the volumetric acquisition at the rate of 20 frames per second with the maximum volume of interest of 315 × 315 × 80 μm^3^. The axial extent of point-spread-function (PSF) was 17.6 ± 1.6 μm and 90.4 ± 2.1 μm with the ETL operating in either stationary or resonant mode, respectively, revealing significant depth elongation by the resonant mode ETL microscopy. We further demonstrated the utilities of the ETL system by volume imaging of cleared mouse brain *ex vivo* samples and *in vivo* brains. The current study foregrounds the successful application of resonant ETL for constructing a basis for a high-performance 3D line-scan confocal microscopy system, which will enhance our understanding of various dynamic biological processes.

## 1. Introduction

For the mechanistic dissection of operational principles of biological phenomena, visualization of dynamic cellular activities *in vivo* is of great importance [1]. Compared to other approaches, real-time imaging of the biological structure and function has only been feasible by optical approaches [2, 3]. Among currently available optical imaging modalities, confocal laser scanning microscopy offers several benefits, such as optical sectioning for 3D imaging and high resolution [4, 5]. However, a limited number of innovative advances in confocal microscopy have been developed for monitoring rapid and dynamic biological processes within tissues like mammalian brains.

In addition to the traditional confocal volumetric imaging, which was achieved by moving the sample along z-direction [6], multiple strategies have been employed to enhance the speed of image acquisition: 1) rapid axial scanning of multiple focal planes using an electrically tunable lens (ETL)[7, 8], 2) utilizing mechanical devices such as deformable mirrors [9], spatial light modulator [10], and adaptive lens [11]. Although these modalities can achieve 3D images at relatively high resolution, the axial scanning rate limits the speed of volumetric imaging. Although alternative strategies including confocal microscopy with multiple pinholes/slits/prisms [12–14] which enables volumetric imaging at a video rate in a relatively large field of view, these methodologies suffer from a limited number of layers detected in the axial direction as well as relatively low optical resolution.

Here, we developed a microscopy technique that enables projected 3D images of an entire volume at a video rate with high optical resolution without suffering constraints in the number of acquired layers. By combining the working principle of a resonant ETL, we named this system an elongated line-scan confocal microscopy (eLSCM). The line-scan confocal microscopy is performed by the addition of a cylindrical lens along with one axis in a confocal point scanning path to achieve 2D imaging at the hundred frames per second. By taking advantage of line-scan confocal microscopy, we conjugated this system with the ETL for vertically averaging each line scan. Conventional confocal microscopy with an ETL had a limited speed in image acquisition due to its 3-axis scanning: two in lateral and one in the axial direction. In contrast, our eLSCM has only one lateral scanning with a fast-axial averaging, which can boost up the speed of volumetric imaging. We demonstrate the utilities of our eLSCM by imaging fluorescently labeled neurons *ex vivo* and cerebral blood vessels *in vivo*.

## 2. Methods

### 2.1. Optical implementation

The schematic diagram of the eLSCM is shown in Fig. 1a. Briefly, in the illumination path, a 475nm laser (Changchun New Industries Optoelectronics Technology Co., Ltd., China, MBL-III-473-100mW) was used as an excitation source. The laser beam size was magnified by three times using a 4-f system (f = 50 mm and f = 150 mm) before it propagated into a cylindrical lens (f_cyl_ = 50 mm), which generated a line pattern parallel to the x-axis at the image plane (side view). The cylindrical lens was shown in the front view as well as a side view (Fig. 1a). A one-axis scanning mirror (Thorlabs, GVS002) was located right after the cylindrical lens to perform as the sweeping light source in a range of 2-degree angle. This laser swept parallel to the y-axis (front view) as shown in Fig. 1b to combine the line pattern from the cylindrical lens for 2D scanning. The elliptical beam propagated to the scan lens (f_scan_= 60 mm) and the first tube lens (f_TL1_ = 100 mm) (Fig.1a). Given that the objective lens was predetermined with the specific numerical aperture value, which was directly correlated with the resolution and the field of view [15], the beam-sheet was connected with the other 4-f system to balance these limitations. This 4-f system consisted of the ETL (Optotune, EL-10-30-C-VIS-LD, generated at 140mA) and the second tube lens (f_TL2_= 150 mm) to fill the light at the back focal plane of the objective lens (Olympus, 20X 0.5 NA, water immersion, UMPLFLN20XW). A dichroic mirror (Semrock, FF552-Di02-25×36) was placed between the first tube lens and the 4-f system of the ETL, to perform fast axial scanning, and to guide the detection path followed by the scan path (Fig. 1b). The resonant frequency of the axial scanning using the ETL (~400 Hz) was 20 times faster than the speed of the lateral image scanning (~20 Hz). Therefore, the axially averaged lines were sufficient for generating a projected 3D image with the elongated PSF in the line scan mode. We further evaluated the effect of the ETL-based 4-f system on the resolution (Supplementary Fig. S1).

**Figure 1.**
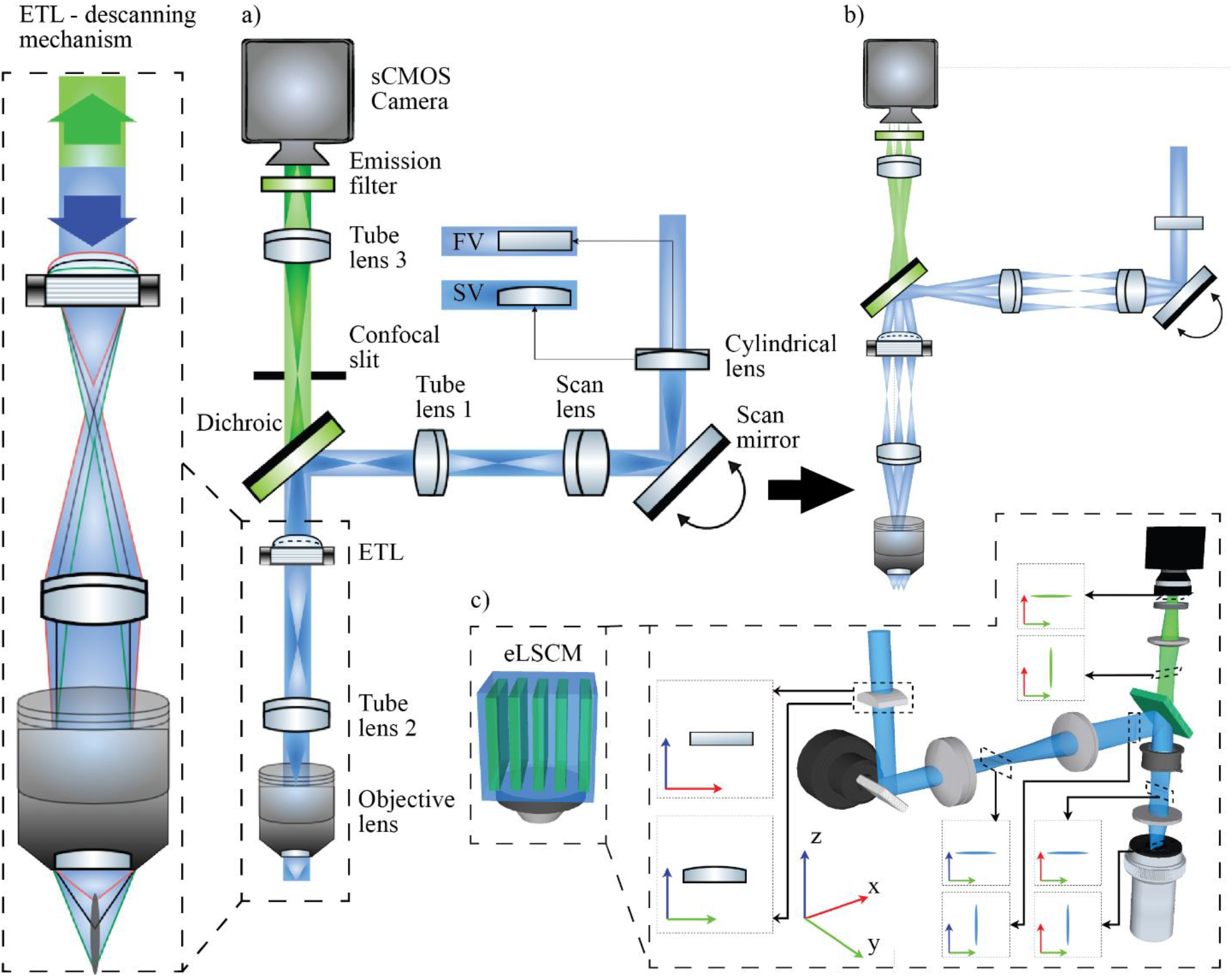
The schematic diagram for ETL-based eLSCM where a) the working principle of the ETL-based eLSCM in front view and side view based on the cylindrical lens, b) the scanning view and direction of the scan mirror on the system, c) the 3D view of the system. FV: front view, SV: side view.

In the detection path, the emitted fluorescent signal propagates back to the tube lens #2 and the ETL, enabling an “ETL descanning mechanism”. We placed a variable optical slit (VA100C/M, Thorlabs) to block the out-of-focus fluorescence signal from the sample. A plano-convex lens (f_TL3_= 60mm) was placed between the dichroic mirror and a detector to generate a line pattern at the detector surface. In our system, an sCMOS camera (PCO.edge 4.2 mono, PCO imaging) was used in a rolling shutter mode to capture the full image of 360 × 360 pixels at 20 fps. All the camera components and devices were driven by open-source software (μ-Manager 1.4) [16]. The hypothetical XYZ-fluorescent signal through the slit and other ideal beam shapes were illustrated in Fig. 1c.

### 2.2. PSF measurement with fluorescent beads

Spatial resolution was measured using fluorescent beads (0.5 μm, Thermo Scientific, G500) [17]. The sample holder (Thorlabs, MAX3SLH) was mounted on a translation stage (Sutter Instrument, MPC-200). The ETL is generated at stationary mode and acquired the image of the bead at continuous layers with the step size is Δz = 0.5 μm. The full volume images of the bead are then reconstructed by the 3D viewer function in ImageJ for images in z-direction [18]. We measured the intensity cross-section profile of each fluorescent bead image in either XY or XZ direction. The intensity data from each bead image was normalized to calculate the full width at half maximum.

### 2.3. Ex vivo imaging of neurons in a cleared brain slice

A coronal section of a cleared brain was prepared from a Thy1-YFP mouse [19]. The experiment was performed in two different modes. First, the translation stage moved the sample toward the objective lens with a step size of Δz = 1.0 μm to acquire full volumetric imaging with the stationary ETL. Second, the image of the cleared brain in the same region was acquired in the resonant ETL mode without moving the translation stage. To visualize the volumetric stack in a 2D image, we used the Time-Lapse Color-Coded Plugins of ImageJ software.

### 2.4. In vivo imaging of blood vessels in a mouse with a cranial window

An open skull craniotomy with a diameter of 5 mm was prepared using C57BL/6 mice. We placed a circular cover glass with a diameter of 6 mm on the open skull and sealed it with dental cement. We used Zoletil/Xylazine mixed in saline solution for anesthesia with the dose of 60/10 mg/kg body weight. The body temperature of an animal was kept stable at 37.0–37.5°C. For *in vivo* blood vessels, imaging was also performed in two different imaging modes. First, after identifying target blood vessels, the volumetric blood vessel images were taken with the step size of Δz = 1.00 μm with a stationary ETL mode. Next, the same area was imaged using a resonant ETL mode for 7.5 seconds at 20 volumes per second. During the post-processing, we used the Time-Lapse Color-Coded Plugins of ImageJ software to highlight the relative depth of blood vessels.

## 3. Result

### 3.1. The ETL characterization with the driving frequency

An ETL can change the focal length based on the driving current [20]. When a shift is applied between two current states, the ETL requires a certain settling time for its stabilization, which limits the ETL’s maximum rate of synchronization during the imaging [21]. As each ETL has its unique settling time and damping property [21], we validated the effect of frequency generation on the scanning range of our system in Figure 2, where the settling frequency was defined as:

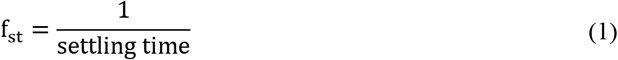

**Figure 2.**
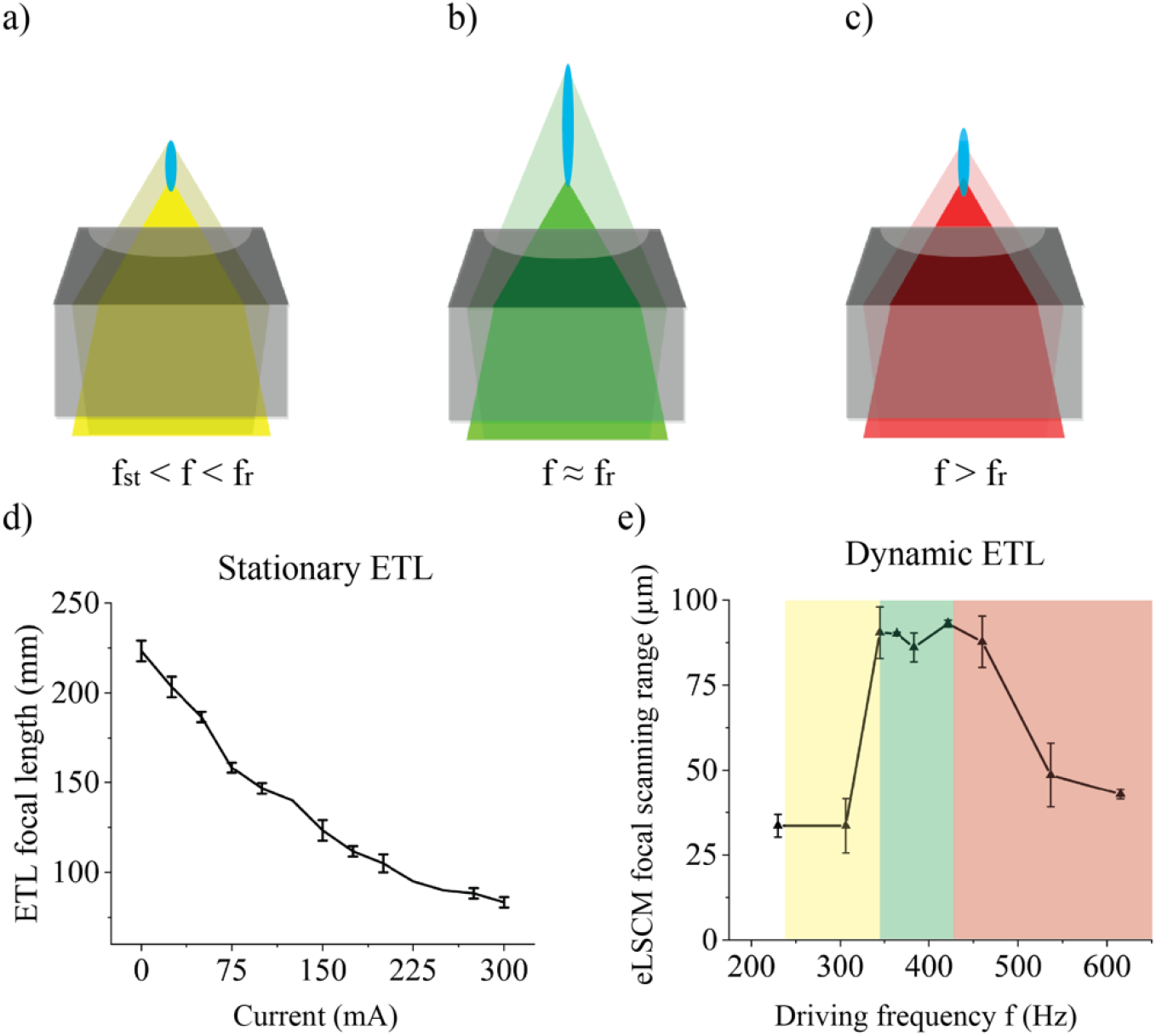
The effect of driving frequency on the axial ETL scanning range. When the driving frequency of ETL is a) greater than settling frequency but lower than resonant frequency, b) approximately resonant frequency, c) greater than the resonant frequency, the elongation of the point-spread function varies (noted in blue). d) the stationary ETL focal length decreases with increasing applied current, e) the dynamic ETL focal scanning range depends on the driving frequency of ETL when the driving current was I(t) = I_0_ sin(2πft) + I_st_. (f_st_ settling frequency, f: driving frequency of ETL, f_r_: resonant frequency, I0: dynamic current amplitude in ETL, I_st_: current at the stable stage of ETL), e) the frequency respond f in scanning range of eLSCM system when f > 0

Theoretically, if the driving frequency of the ETL is lower than the settling frequency, the scanning range will be dependent on the applied current and frequency. However, the current ETL f_st_ was 60 Hz, which was too slow to combine with the lateral scanning for video-rate eLSCM imaging. If the modulation frequency is higher than the settling frequency of the ETL but is lower than the resonant frequency, as in Fig. 2a, the ETL will be over-damped and cannot achieve the desired elongation of a focal length. Therefore, the scanning range will be determined by the ETL driving frequency. At the resonant frequency, the ETL scanning range is maximized (Fig. 2b). In contrast, when the ETL is driven at a higher than the resonance frequency, it is over-damped, resulting in a reduced scanning range (Fig. 2c).

We first calibrated the ETL focal length with respect to the driving current. When the frequency was set at 0 Hz (stationary ETL), the focal position was identified and recorded at each driving current. For example, the focal length was ~ 130 mm with a current of 140 mA. When a higher current was applied in the range 0 – 300 mA, the focal length got shorter from 230 mm – 90 mm shown in Fig. 2d. The focal scanning range varied depending on the driving frequency when the frequency was non-zero (dynamic ETL). At the resonant frequency of the ETL, we set the current in the range between 110 mA and 170 mA, which corresponds to the focal length of approximately 120 mm to 150 mm, respectively. This configuration ensures to fill the back aperture of the objective lens via the tube lens with focal length of 150 mm before entering into the objective lens (Fig. 1 left box).

The ETL driving current at resonance can be described as below:

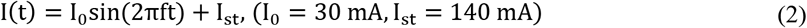

Since the ETL settling frequency of f_st_ = 60 Hz was too slow for fast volumetric imaging with our line-scan confocal microscopy, we measured the focal scanning range in FWHM with respect to the driving frequency from 200 Hz to 600 Hz as shown in Fig. 2e. The maximum scanning range of 90 ~ 100 μm was obtained with the driving frequency range of 340 ~ 420 Hz, and thus we used 400 Hz for our ETL-based eLSCM.

### 3.2. Axially elongated point-spread function measurement

To measure the PSF, we employed two ETL modes to compare resolutions with different modes, i.e., stationary and resonant ETL modes. With a stationary ETL mode with a driving current of 140 mA, the lateral full-width-half-maximum (FWHM) was δx = 2.0 ± 0.5 μm (n = 3) and the axial resolution FWHM was δz = 17.6 ± 1.6 μm (n = 3) (Fig. 3a). With a resonant ETL mode at 400 Hz, a sinusoidal current from 110 mA to 170 mA was used with corresponding focal lengths of 120 mm to 150 mm, respectively. The lateral FWHM was at δx = 3.2 ± 0.3 μm (n = 3) and the axial FWHM was elongated by 5.1 times to δz = 90.4 ± 2.1 μm (n = 3) (Fig. 3b). We acknowledged that the sinusoidal nature of driving current of an ETL might result in higher intensity values at two peaks due to transient slowdown. While brighter at the top position, this phenomenon might be beneficial for imaging a scattering medium due an elevated signal at deeper layers. On the other hand, the lateral resolution remained approximately equivalent within one-micron range between two modes (Supplementary Fig. S2).

**Figure 3.**
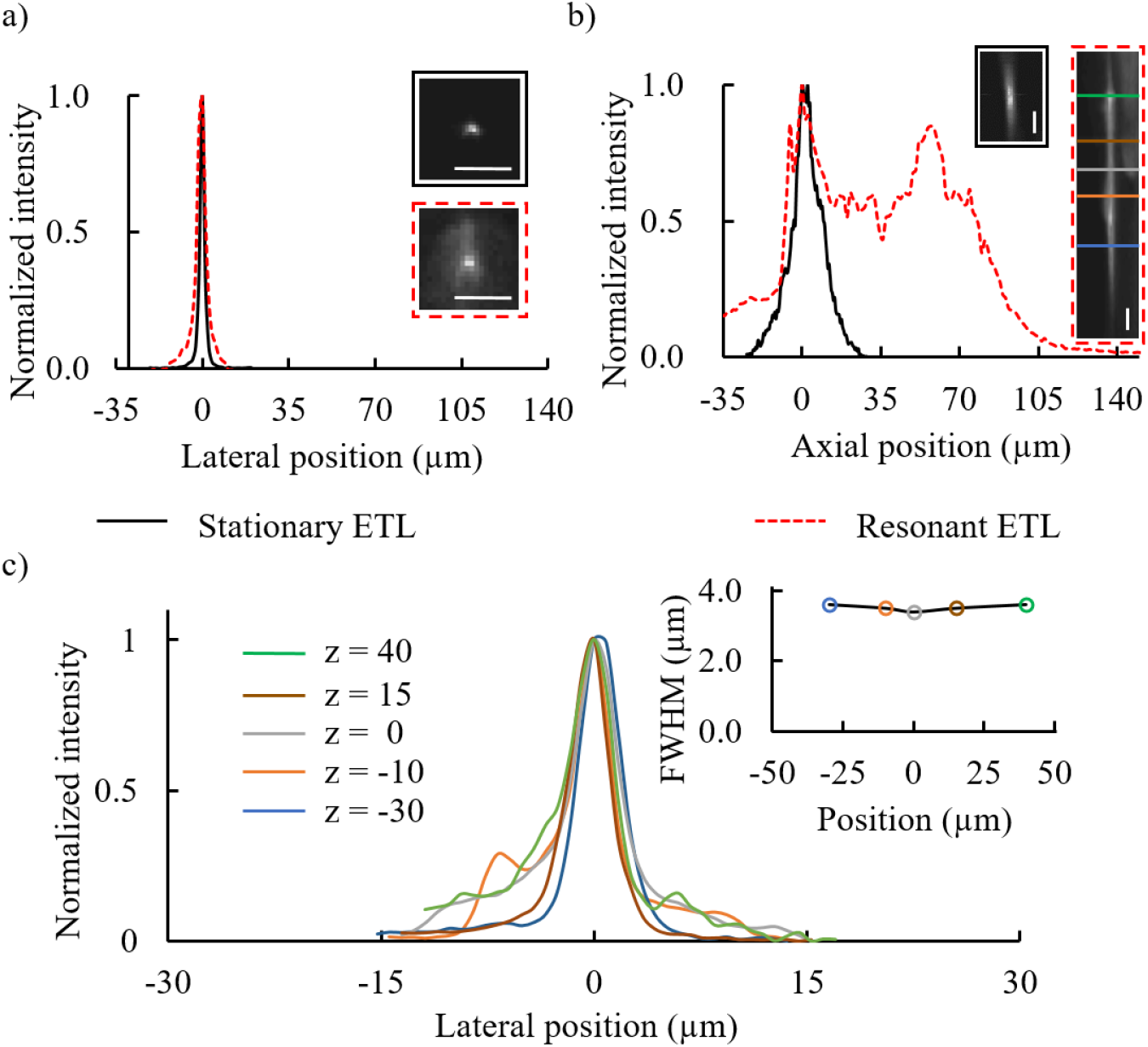
PSF measurement in XY and XZ transverse direction with 0.5 μm fluorescent beads, a) the lateral FWHM value, b) the axial FWHM value of the system when ETL is at the stationary stage at 140 mA and resonant stage from 110 mA to 170 mA at ~400 Hz and c) the profile of lateral PSF at the chosen five layers in the axial direction at resonant ETL stage. Scale bar: 10 μm

The FWHM data at different depths were shown in Fig. 3c, where each color represented the corresponding plane in the axial direction. We observed that the most significant change in normalized intensity in the axial direction was approximately 6%, indicating that the elongated PSF’s lateral resolution remained consistent throughout different depths.

### 3.3. Volumetric imaging of cleared brain ex vivo

After PSF characterization, we imaged a cleared brain section harvested from a Thy1-YFP transgenic mouse (Fig. 4a). Cortical regions of the brain were imaged using our ETL-based eLSCM system. The experiment was performed in two different modes: stationary and resonant ETL with corresponding driving current I(t). In the stationary ETL mode, we imaged neurons at each layer with the step size of Δz = 1 μm with the total range of 80 μm. Fig. 4b showed the result at the middle plane of the brain. Next, multi-layer z-stack images were shown in Fig. 4c, where the color illustrates the axial positions of the layer. In the resonant ETL mode, identical cortical neurons were visualized at the central position of z = 0 μm FOV at 20 fps averaging for 5 seconds covering a volume of 315 × 315 × 80 μm^3^. (Supplementary V1).

**Figure 4.**
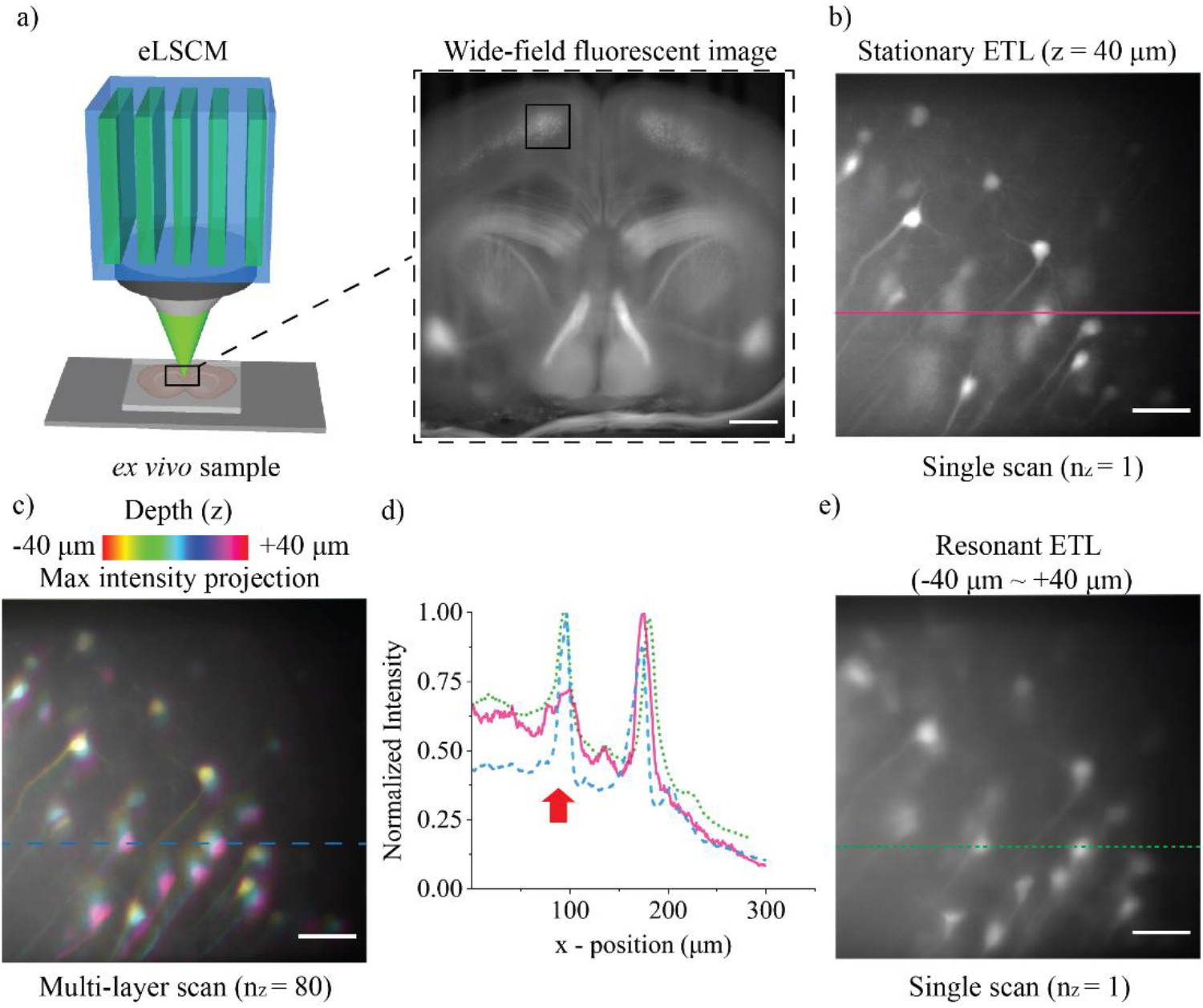
Volumetric imaging of a cleared brain slice *ex vivo* a Thy1-YFP brain. a) wide-field fluorescence image, scale bar: 500 μm, (b-c) cortical neurons imaged with stationary ETL mode with the step size Δz = 1 μm, total ~80 layers at −40 μm-layer and projection image, respectively, e) cortical neurons imaged with resonant ETL mode with an extended depth of field, d) the normalized intensity plot profile of the dotted line in different modes b), c) and e). Scale bar: 50 μm.

To demonstrate the elongated PSF’s effect, we performed a single scan of the ETL in a resonance mode, as shown in Fig. 4e. The intensity profiles of the neuronal image located along the identical line were shown in three modes: single scan in a resonant ETL, single and multi-layer scan in the stationary ETL mode in Fig. 4d. While the normalized intensities of single scan in resonant ETL and multi-layer scan in stationary ETL were closely matched, the data plotted in single scan of the stationary ETL mode indicated discrepancy (arrow), show the loss of signal from a neuron located out of focus within the brain cortex. This demonstrated that the ETL-based eLSCM single frame image contains the average signal of a total 80 μm-depth of the sample. However, due to the limitation in the speed of the ETL, the background noise in the acquired image at the resonant ETL mode was higher than that from the multi-layer scan image with the stationary ETL mode.

### 3.4. Volumetric imaging of blood vessels of a mouse brain in vivo

For *in vivo* experiment, we performed craniotomy to place an optical window [22]. Under anesthesia, the animal was injected with fluorescein isothiocyanate – Dextran 500 kDa-Conjugate (FITC) via retro-orbital injection [23, 24].

FITC-labeled cerebral vessels were visualized with the eLSCM, as shown in Fig. 5a. The eLSCM was employed in two modes. In the stationary ETL mode, we imaged blood vessels in multiple 2D planes using a translation stage; each plane represented one image at 20 fps with the FOV of 315 × 315 μm^2^. Next, we captured the image of one selected layer (Fig. 5b) prior to 3D imaging with an axial range of 80 μm (Fig. 5c). With the resonant ETL mode, we imaged blood vessels in the same FOV (Fig. 5e); the full videos of a 3D stacked image and the ETL-based eLSCM of cerebral blood vessels are shown in Supplementary V2.

**Figure 5.**
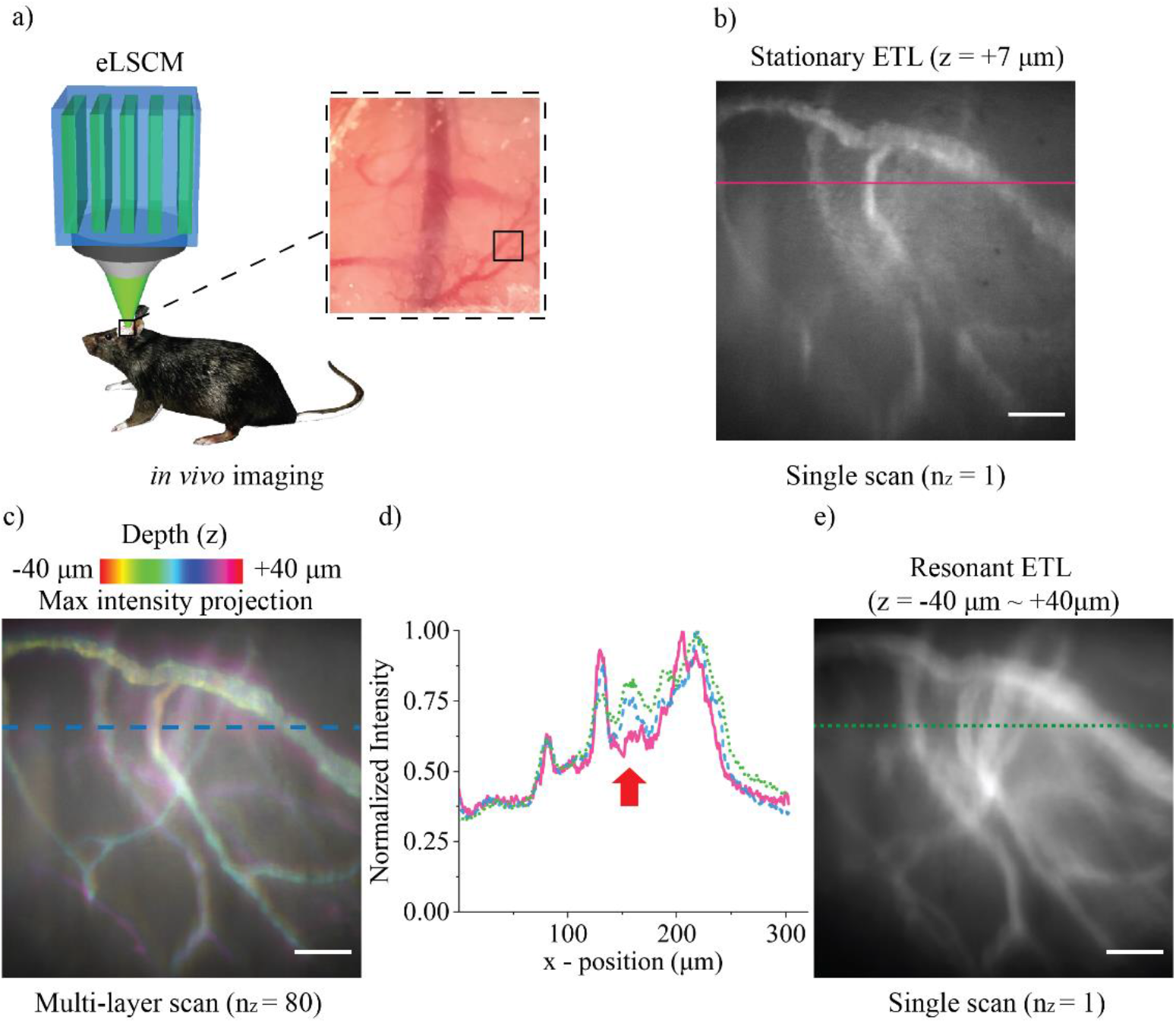
Imaging blood vessels in the mouse brain *in vivo*. a) the schematic diagram of *in vivo* brain imaging experiment, b) the blood vessels imaging of +7 μm-layer in a stationary ETL mode, total ~80 layers, with the step size Δz = 1 μm, c) the total volumetric color-coded by depth image of the blood vessels, e) the single scan from video-rate imaging extended depth of field of blood vessels in resonant ETL mode for 7.5 seconds, and d) the normalized intensity cross-section of two imaging modes (pink line: cross-section of single scan stationary, blue line: cross-section of volumetric in stationary, green line: cross-section of extended depth of field in the resonant stage). Scale bar: 50 μm.

Line intensity profiles were taken from the same location in 3 different modes of our system. At the stationary stage, the cross-section intensity profiles were obtained from the single-layer scan (red line) and the max intensity projection of the multi-layer scan (blue dotted line). At the resonant stage, the line intensity profile was taken from an image in a total of 7.5 seconds scan (green dotted line). All intensity profiles were normalized to highlight differences between images acquired from two different modes (Fig 5e). The peak intensity levels were relatively well-matched when we compared the result from the volumetric multi-layer scan of the stationary stage to a single scan in the resonant ETL mode. In contrast to this, by comparing the single scan in both stationary and resonant mode, indicating the loss of structural and functional information of blood vessels in stationary stage and informational completeness of blood vessels visualization in resonant mode at video-rate.

## 4. Discussion

We developed an extended depth-of-field, line-scan confocal microscopy by utilizing a resonant ETL to elongate the PSF in the axial direction by 5.1 times to cover approximately 90.4 μm in depth. Compared to the system where the ETL was attached beneath the objective lens (Supplementary Fig. S1), the lateral optical resolution was 1.5 ± 0.1 μm, which was slightly better than the ETL-based eLSCM (the lateral optical resolution: 2.0 ± 0.5 μm) (Fig. S2b). However, in a resonant ETL mode, the lateral optical resolution in an ETL-based eLSCM was 3.2 ± 0.3 μm (Fig. S2c). In addition, the results obtained from the line scan confocal microscopy without an ETL showed that the lateral and axial resolutions obtained were 0.8 ± 0.01 μm and 6.3 ± 0.1 μm, respectively, better than those of stationary ETL mode (Fig. S2c). The degradation in lateral and axial optical resolution in a stationary ETL mode is likely caused by the aberration of ETL and the blurring effect caused by the stability of driving current the ETL, affecting the resultant point spread function measurement.

By using an *ex vivo* brain slice, we demonstrated that our eLSCM system could image at the maximum z = 80 μm in the full volumetric dimensions of 315 × 315 × 80 μm^3^ at 20 fps. We also imaged cerebral blood vessels *in vivo* towards measuring blood flow velocity [25–27]. We expect that our eLSCM system can perform other *in vivo* experiments such as volumetric calcium imaging of sparsely labelled cortical neurons using fluorescent dyes or genetically encoded calcium indicators [13, 14] to correlate neural activities during animal behaviors [28, 29]. However, we acknowledge that the current approach can only be applied to a sparsely labeled target as the system provides an averaged axial projection image.

Other microscopy techniques utilizing the elongated PSF for volumetric imaging suffer from compromised lateral and axial resolution [13], or limited axial detection range [30], even the aberration caused by the phase mask along with the pinhole which not compatible slits [31–33]. These aberrations could be reduced by adaptive optics [34, 35]. The major caveat of our system is that the empirically determined resonant frequency sets the speed of image acquisition. This limit can be resolved by employing a TAG lens [36–38] or remote focusing [39]. In support of this notion, the resonance frequency of a commercially available TAG lens is at 70 kHz (Mitutoyo), which can enhance the acquisition speed of our eLSCM to hundreds of frames per second. If we can synchronize the scanner signal with a high-speed axial scanning unit, a high resolution in the multi-plane imaging method might be achievable [40].

Taken together, we developed novel microscopy by combining line-scan confocal microscopy and a resonant ETL-based axial scanning unit to elongate PSF enabling volumetric projection imaging at video rate towards functional imaging of dynamic biological processes.

## 5. Acknowledgments

## 5.2. Acknowledgments

The work was supported by The GIST Research Institute (GRI) research collaboration grant funded by GIST in 2021 and the 2021 Joint Research Project of Institutes of Science and Technology, a grant from the National Research Foundation of Korea (N.R.F.) funded by the Korean government (MEST) (NRF-2019R1A2C2086003), the Brain Research Program through the N.R.F. funded by the Ministry of Science, I.C.T. & Future Planning (NRF-2017M3C7A1044964), and the Korea Medical Device Development Fund grant funded by the Korea government, the Ministry of Science and ICT, the Ministry of Trade, Industry and Energy, the Ministry of Health & Welfare, the Ministry of Food and Drug Safety (Project Number: 1711138096, KMDF_PR_220200901_0076) to EC.

HSJ is supported by Singapore National Medical Research Council Research Grant (NMRC/OFIRG/0050/2017) and Singapore National Science Foundation Research Grant (NRF-CRP17-2017-04).

ETJ is supported by the National Research Foundation of Korea(NRF) grant funded by the Korea government(MSIT) (No. NRF-2021R1A5A1032937).

The authors thank Mr. YoungSeung Yo for helping in preparing sample for this works. We also thank Dr. Mohiuddin Khan Shourav for helping in figure panel preparation.

