## Supplementary S1 and S2 for "Fast volumetric imaging with line-scan confocal microscopy by an electro-tunable lens"

### Supplementary material

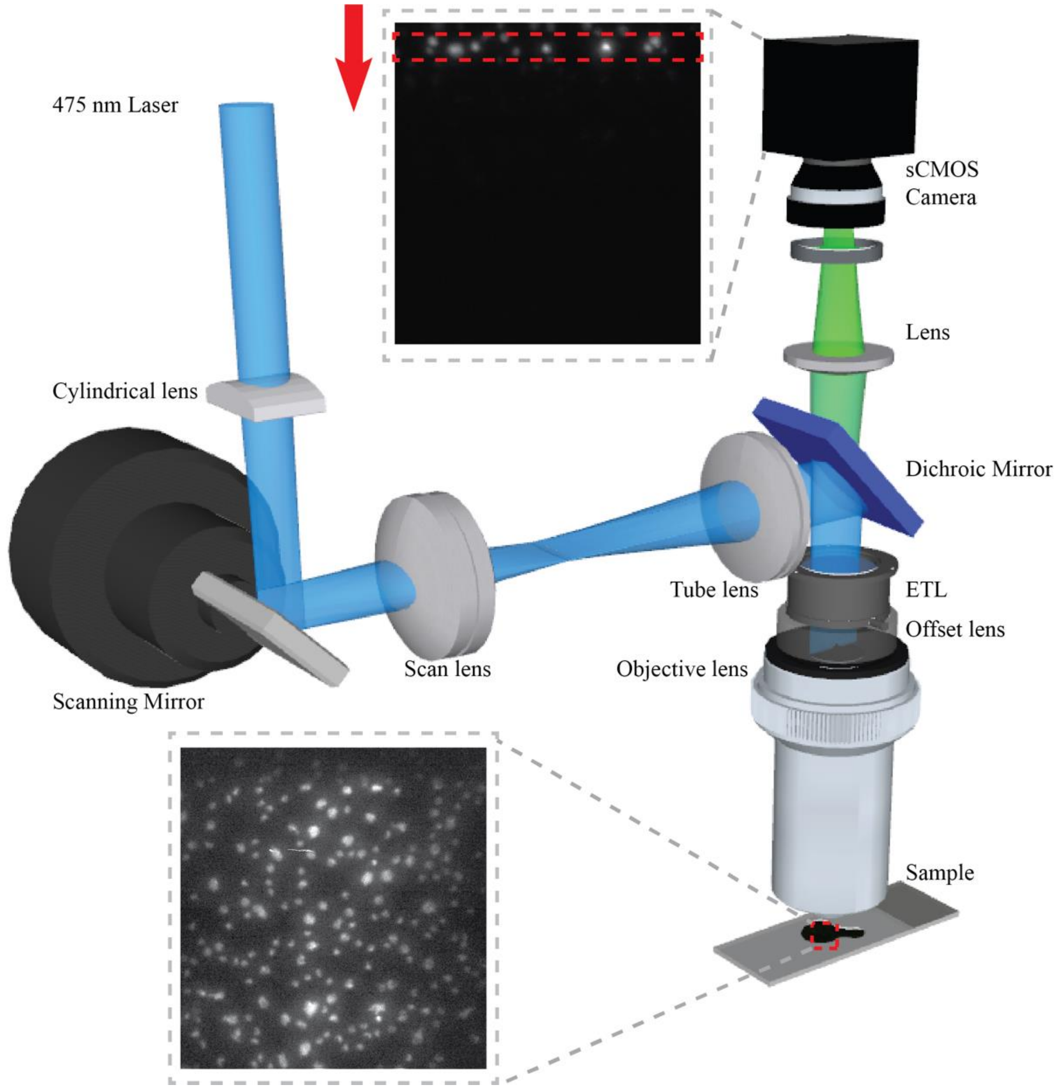


**Figure S1. The schematic of ETL-based eLSCM while ETL is constructed above the objective lens with the offset lens (concave lens) instead of a 4-f system manufacturing.**

The laser beam size is magnified 3 times by a 4-f system (f = 50 mm and f = 150 mm, respectively) before it propagates the cylindrical lens ($f_{cyl}$ = 50mm) to generate the line pattern at the image plane, parallel to the x-axis. Here we use the one-direction scanning mirror (re-modified from Thorlabs GVS002) right after the cylindrical lens to perform the sweeping laser in a range of 4-degree angles, parallels to the y-axis The scanning of the mirror combine the line pattern from the cylindrical lens to perform a full image. The elliptical beam then propagates the Scan lens ($f_{scan}$ = 60 mm) and Tube lens ($f_{TL}$ = 250 mm) to full fill and then conjugated with the back focal plane of the objective. A dichroic mirror (Semrock, FF552-Di02-25x36) and the ETL (Optotune, EL-10-30-C-VIS-LD) system are placed in between the Tube lens and Objective lens (Olympus, 20X 0.5 NA, water immersion, UMPLFLN20XW) to enable the axial scanning and guide the detection path follows by the scan path. We place a plano-concave lens (f = -50mm) to prevent the coma aberration that normally occurs in the imaging system. The fluorescent signal after excitation from the sample propagates the dichroic mirror. An achromatic doublet lens (f = 150mm) is placed in between the dichroic mirror and detector, the position of the lens is conjugated with the back focal plane of the objective lens and the detector surface. An sCMOS camera is used to capture the full image which is swept by the scanner mirror as indicated.

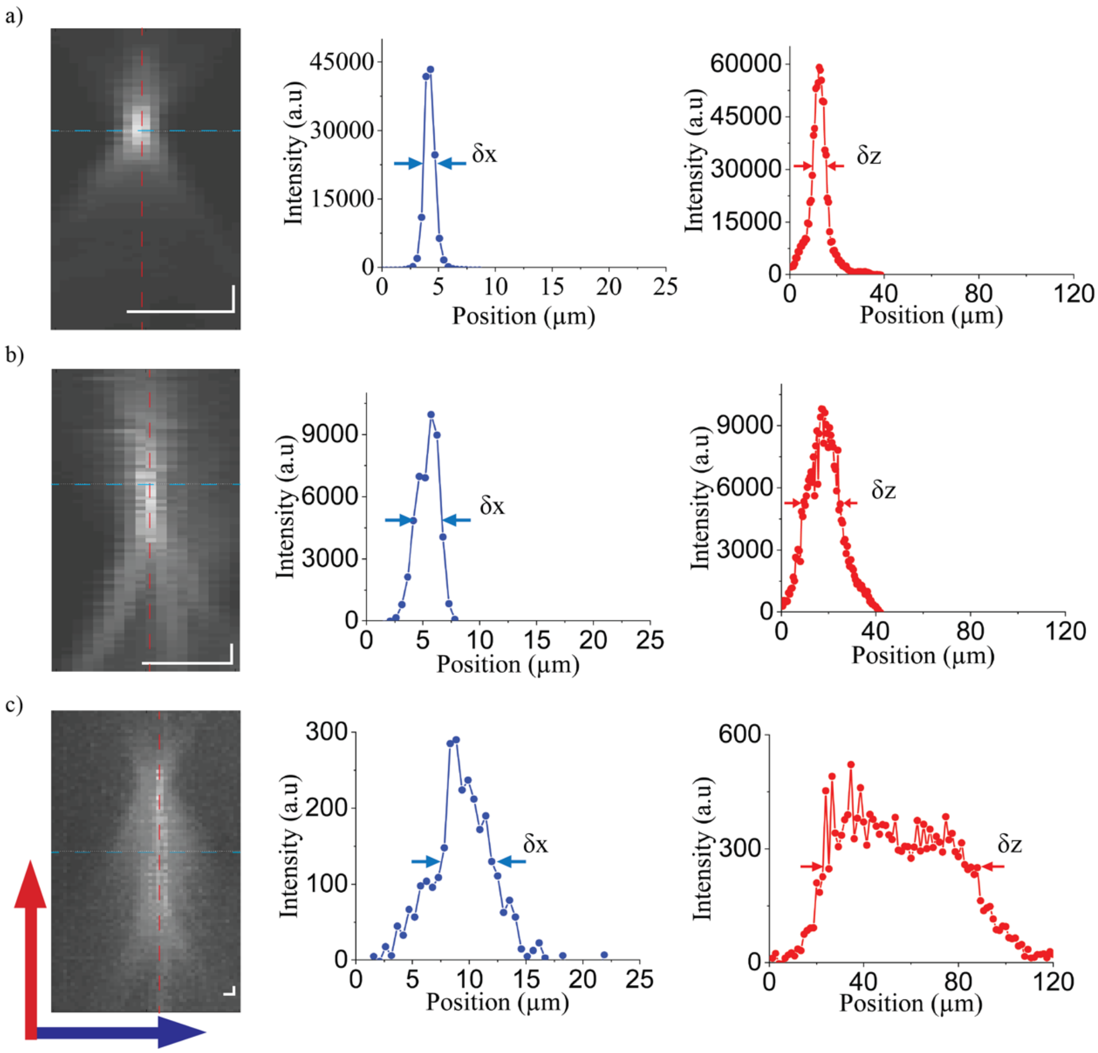


**Figure S2. PSF measure of the system in Figure S1.** Lateral (blue) and axial (red) spread function of ETL-based eLSCM without ETL related 4-f system with 0.5 μm fluorescent bead and its profile where a) ETL and offset lens is removed, b) ETL and offset lens is attached but is not generated and c) ETL is generated in sinusoidal wave function from 0mA to 60mA at around 350 Hz. Scale bar: 5 μm.

We generated the ETL in two different categories to compare the resolution in each stage: without ETL, ETL off, and eLSCM. At the first stage, the lateral full-width-half-maximum (FWHM) achieved is $\delta x = 0.8 \pm 0.01$ μm (n=3) while the axial resolution FWHM is $\delta z = 6.3 \pm 0.1$ μm (n=3) (Fig. S2a). We then attached the ETL in the system, since the ETL is a lens; therefore, the combination of it and objective lens may be degraded, we then got the lateral and axial resolution is $\delta x = 1.5 \pm 0.1$ μm (n=3) and $\delta z = 18.7 \pm 0.8$ μm (n=3), respectively (Figure S2b). The lateral FWHM then is achieved at $\delta x = 5.2 \pm 0.1$ μm (n=3) while the ETL is generating for eLSCM while in the axial direction, the FWHM of value is elongated to $\delta z = 65 \pm 2.2$ μm (n=3) by the effect of axial scanning of the ETL (Figure S2c).
