## Supplementary figures and images for "Fast volumetric imaging with line-scan confocal microscopy by an electro-tunable lens"

### Supplementary V1

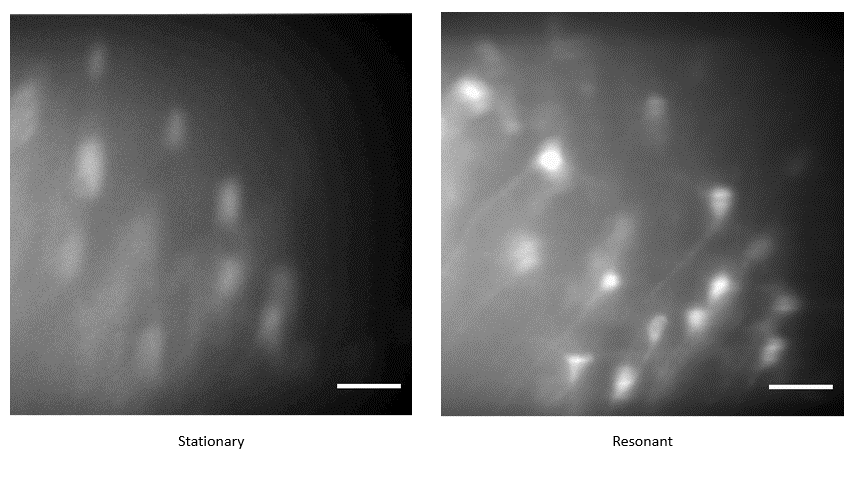

### Supplementary V2

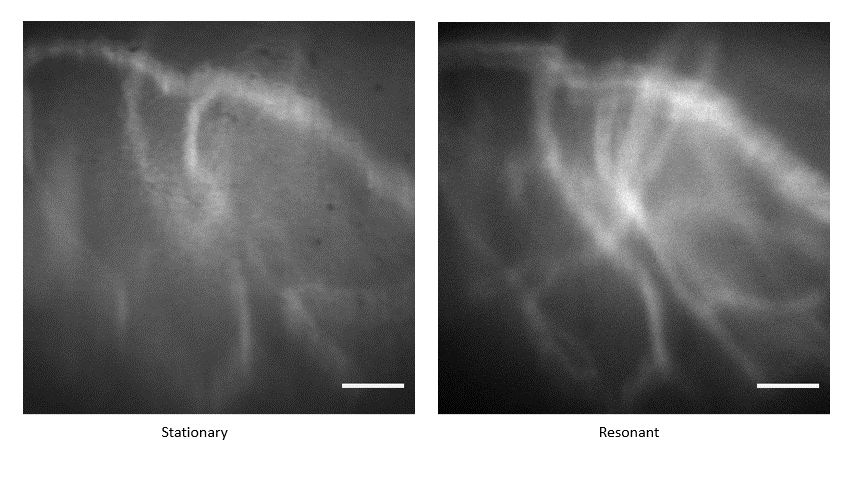
